# Single-cell multimodal profiling of atherosclerosis identifies CD200 as a cell surface lineage marker of vascular smooth muscle cells and their derived cells

**DOI:** 10.1101/2023.09.29.557779

**Authors:** Alexander C. Bashore, Allen Chung, Chinyere Ibikunle, Hanying Yan, Chenyi Xue, Mingyao Li, Robert C. Bauer, Muredach P. Reilly

**Affiliations:** Division of Cardiology, Department of Medicine, Vagelos College of Physicians and Surgeons, Columbia University, New York; Department of Biological Sciences, Columbia University, New York; Department of Biostatistics, Epidemiology and Informatics, University of Pennsylvania Perelman School of Medicine, Philadelphia, PA; Irving Institute for Clinical and Translational Research, Columbia University Irving Medical Center, New York, NY

## Abstract

Vascular smooth muscle cells (VSMCs) play a central role in the development of atherosclerosis due in part to their capability to phenotypically transition into either a protective or harmful state. However, the ability to identify and trace VSMCs and their progeny in vivo is limited due to the lack of well-defined VSMC cell surface markers. Therefore, investigations into VSMC fate must utilize lineage-tracing mouse models, which are time-consuming and challenging to generate and not feasible in humans. Here, we employed CITE-seq to characterize the phenotypic expression of 119 cell surface proteins in mouse atherosclerosis. We found that CD200 is a highly expressed and specific marker of VSMCs, which persists even with phenotypic modulation. We validated our findings using a combination of flow cytometry, qPCR, and immunohistochemistry, all confirming that CD200 can identify and mark VSMCs and their derived cells in early to advanced mouse atherosclerotic lesions. Additionally, we describe a similar expression pattern of CD200 in human coronary and carotid atherosclerosis. Thus, our data support the use of CD200 as a lineage marker for VSMCs and VSMC-derived cells in mouse and human atherosclerosis.

---

Atherosclerotic cardiovascular disease is the leading cause of death worldwide [1]. Vascular smooth muscle cells (VSMCs) play a central role in atherosclerosis due to their capability to phenotypically transition into either a protective or harmful state. However, the ability to identify and trace VSMCs and their progeny in vivo is limited due to the lack of well-defined VSMC surface markers [2]. Current investigations into VSMC fate must utilize lineage-tracing mouse models, which are time-consuming, challenging to generate, and not feasible in humans. Here, we employed CITE-seq to phenotypically characterize the expression of 119 cell surface proteins in mouse atherosclerosis. We also performed CITE-seq in human atherosclerosis and found that CD200 is a highly expressed and specific marker of VSMCs that persists even with phenotypic modulation.

To characterize cell types at different stages of atherosclerosis, we used an LDLr-/-; ROSA26LSL-ZsGreen1/+; Myh11-CreERT2 mice, which permanently induces the expression of ZsGreen1 in VSMCs and their progeny following tamoxifen administration. At four different durations of western diet feeding (0, 8, 16, and 26 weeks), ZsGreen1+ and ZsGreen1-cells were FACS sorted and submitted for CITE-seq profiling **(Fig [A])**. 13 distinct cell clusters were identified, including macrophages, T cells, endothelial cells, VSMCs, and fibroblasts. Based on ZsGreen1 status, VSMC-derived cells mainly comprised the VSMC 1, VSMC 2, and modulated VSMC populations (**Fig [B]**). Our analysis identified CD200 as the most highly expressed and specific marker of VSMCs and VSMC-derived cells (**Fig [C]** and **Fig [D]**). There was also a marked lower and less specific expression on fibroblast and endothelial cell clusters. Additionally, the expression of CD200 was consistent and maintained throughout different durations of atherosclerosis up to and including 26 weeks (**Fig [E]**).

**Figure.**
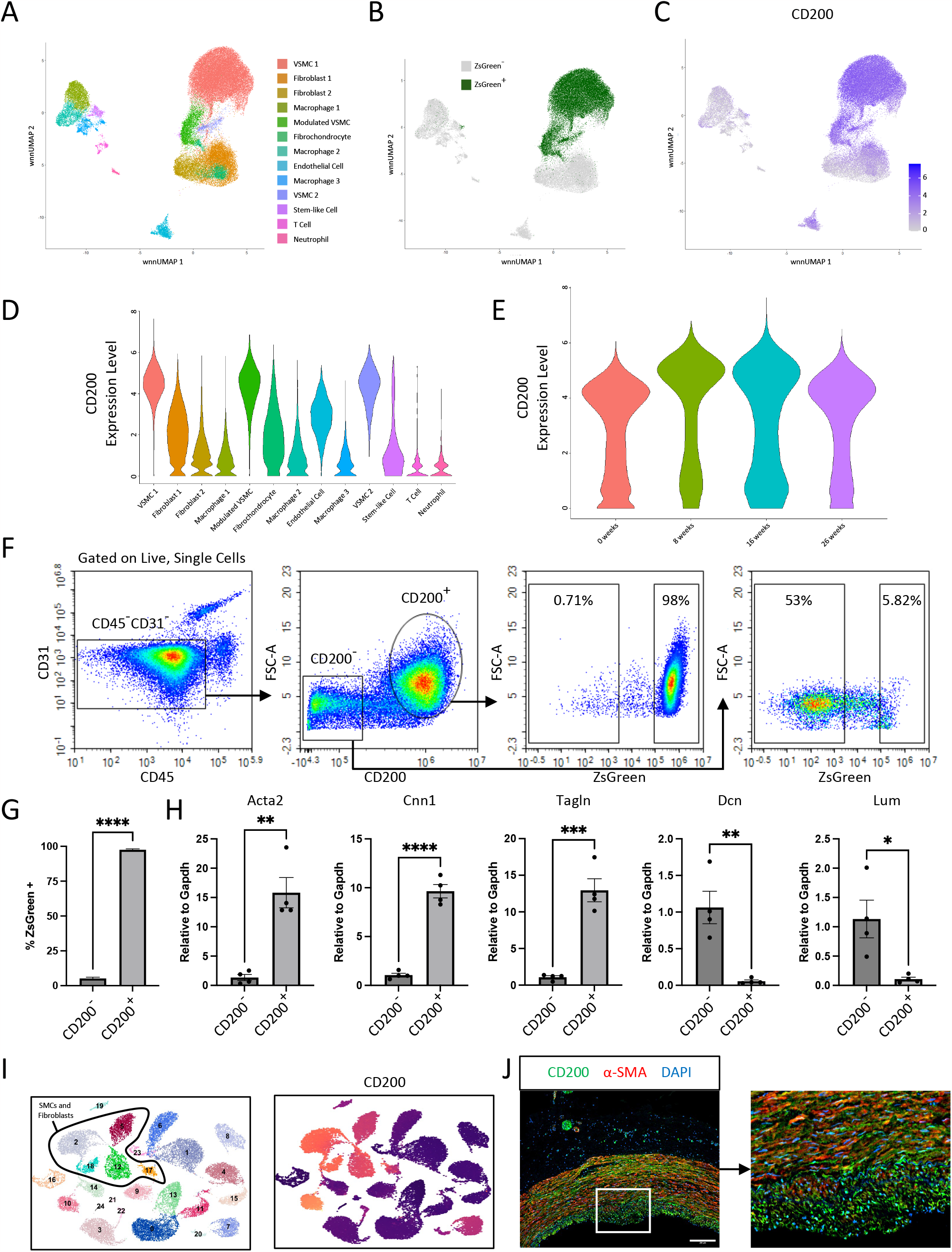
Identification and validation of CD200 as a cell lineage marker in mouse and human atherosclerosis. (A) UMAP visualization of multimodal integration of all CITE-seq data identifying 13 distinct cell populations. (B) UMAP depiction of ZsGreen status. (C) Feature plot showing the expression of CD200 protein across all cell clusters. (D) Violin plot quantifying the expression of CD200 in each cell cluster. (E) Violin plot quantifying the expression of CD200 at each time point of western diet feeding. (F) Flow cytometry gating strategy to assess the percentage of CD200+ cells that are ZsGreen+. (G) Quantification of the percent of CD200+ cells that are ZsGreen+. Values are shown as mean±SD, n=2. (H) Relative mRNA expression of Acta2, Cnn1, Tagln, Dcn, and Lum in FACS-sorted CD200+ and CD200-cells. Values are shown as mean±SD, n=4. (I) UMAP visualization of human carotid atherosclerosis CITE-seq data with expression of CD200 across all cells, n=6. (J) Immunohistochemistry staining of human coronary artery sections. CD200 (green), α-SMA (red), and DAPI (blue).

To validate our CITE-seq findings, we developed a flow cytometry panel to determine if CD200 can discriminate between VSMC and fibroblasts and used this panel to analyze cells from the ZsGreen1+/- lineage traced mice (**Fig [F]**). To remove leukocytes and endothelial cells, we excluded CD45+ and CD31+ cells, respectively, so that the remaining cells include all VSMCs and fibroblasts. We next examined the expression of CD200, identifying two distinct populations of CD200+ and CD200-cells. Finally, we inspected the proportion of each population that was ZsGreen1 positive and negative. We discovered that the CD200+ cells were over 97% ZsGreen1+, whereas the CD200-cells were only 5% ZsGreen1+. (**Fig [G]**). For further validation, we sorted CD200+ and CD200-cells that were also CD45-CD31-from C57BL6J mice and assessed the expression of VSMC- and fibroblast-specific genes (**Fig [H]**). The sorted CD200+ cells displayed a 10-15-fold increase in the expression of the VSMC-specific genes Acta2, Cnn1, and Tagln, and a greater than 90% lower expression of fibroblast-specific genes Dcn and Lum compared to CD200-cells. These data support that CD45-, CD31-, CD200+ cells can identify VSMC and VSMC-derived cells extremely effectively and that the expression of CD200 is maintained on VSMCs and VSMC-derived cells throughout atherosclerosis. In support of clinical relevance and translational utility, we corroborated these findings in human atherosclerosis with CITE-seq data from a previously reported carotid endarterectomy dataset and observed a similar pattern (**Fig [I]**) [3]. We also performed immunohistochemistry on human coronary arteries to localize CD200-expressing cells within atherosclerotic lesions. We stained these tissues with α-SMA (to identify VSMCs) and CD200 (**Fig [J]**). There was a high degree of colocalization in the media of the artery with a lack of staining within the adventitia. Within lesion neointima, there was strong CD200 staining; however, there was less colocalization, likely because as VSMCs phenotypically modulate, they lose the expression of contractile VSMC proteins, including ACTA2 [4].

Here, our CITE-seq analysis revealed that CD200 is highly expressed in VSMCs, and this expression persists in their derived cells in the development and progression of atherosclerosis. Although CD200 is known to be produced by many cells, including epithelial, hematopoietic, and endothelial cells [5], ours is the first study to report that CD200 is a marker of VSMCs and their modulated progeny. Based on our VSMC lineage-traced mouse data, leukocytes and endothelial cells are the only other major cell types of the vasculature that express CD200. Therefore, combining CD200 positivity with CD31 and CD45 negativity makes it possible to separate VSMCs from endothelial cells and leukocytes.

Current studies of VSMC origin in atherosclerosis necessitate breeding lineage tracing systems as VSMCs are plastic and can lose expression of canonical contractile genes. Our discovery that CD200 is a persistent VSMC marker, even during VSMC phenotypic switching, abrogates the need to include a lineage-tracing reporter and facilitates investigations of VSMC and VSMC-derived cell roles in atherosclerosis, including critically in humans where VSMC lineage-tracing has limited feasibility. In conclusion, by identifying a cell surface marker of VSMCs, multiple techniques, including flow cytometry, can now be employed to characterize VSMCs and their phenotypically modulated cell types in atherosclerosis and its complications.

